# Nearly Neutral Evolution Across the *Drosophila melanogaster* Genome

**DOI:** 10.1101/212779

**Authors:** David Castellano, Jennifer James, Adam Eyre-Walker

## Abstract

Under the nearly neutral theory of molecular evolution the proportion of effectively neutral mutations is expected to depend upon the effective population size (*N_e_*). Here we investigate whether this is the case across the genome of *Drosophila melanogaster* using polymorphism data from North American and African lines. We show that the ratio of the number of non-synonymous and synonymous polymorphisms is negatively correlated to the number of synonymous polymorphisms, even when the non-independence is accounted for. The relationship is such that the proportion of effectively neutral non-synonymous mutations increases by ~45% as *N_e_* is halved. However, we also show that this relationship is steeper than expected from an independent estimate of the distribution of fitness effects from the site frequency spectrum. We investigate a number of potential explanations for this and show, using simulation, that this is consistent with a model of genetic hitch-hiking: genetic hitch-hiking depresses diversity at neutral and weakly selected sites, but has little effect on the diversity of strongly selected sites.

## Introduction

In 2015, Tomoko Ohta was awarded the Crafoord prize, along with Richard Lewontin, for her contributions to evolutionary biology and population genetics. Among her many contributions, she is best known for her nearly neutral theory of molecular evolution in which she proposed that there is a class of mutations that are substantially influenced by both random genetic drift and natural selection (Ohta and Kimura 1971; Ohta 1972b, 1973, 1977, 1992). In species with large effective population sizes selection is effective and in species with small effective population sizes random drift dominates the fate of this class of mutations. As a consequence, the proportion of mutations that are effectively neutral varies with effective population size. The theory has had a profound influence on how we think about molecular evolution and processes such as the rate of evolution (Lanfear, et al. 2014), the evolution of the mutation rate (Lynch 2010; Lynch, et al. 2016) and genome size (Lynch and Conery 2003).

There are a number of lines of evidence suggesting that there is a class of mutations that are nearly neutral, the vast majority of which are thought to be slightly deleterious. First, putatively functional mutations segregate at lower frequencies in the population than putatively neutral mutations; for example, in many species non-synonymous mutations segregate at lower frequencies than synonymous mutations (Akashi 1999; Cargill, et al. 1999; Hughes 2005), and it has also been shown that mutations in conserved non-coding sequences segregate at lower frequencies than those in neighbouring sequences (Andolfatto 2005; Drake, et al. 2006; Asthana, et al. 2007). Second, the rate of non-synonymous to synonymous substitution is higher along lineages leading to species with small effective population size (note that in most cases the effective population is not directly measured but a surrogate measure, such as body size, is used instead) (Ohta 1993, 1995; Moran 1996; Woolfit and Bromham 2003, 2005; Popadin, et al. 2007). In fact, Ohta produced the very first evidence for her theory by noting that the rate of divergence in protein coding sequences relative to the overall rate of DNA divergence, as measured by DNA-DNA hybridisation, was positively correlated to generation time, a proxy for population size (Ohta 1972a). Third, the ratio of non-synonymous or synonymous polymorphism appears to be greater in those species with smaller effective population size, as measured by synonymous diversity (Piganeau and Eyre-Walker 2009; Elyashiv, et al. 2010; Galtier 2016; Chen, et al. 2017; James, et al. 2017).

Ohta used her theory to explain how the rate of protein evolution could be relatively constant even if the mutation rate varied between taxa; she reasoned that species with large population sizes might have high mutation rates per year, but this would be offset by having a small proportion of effectively neutral mutations (Ohta and Kimura 1971; Ohta 1972b, 1992). However, despite all the qualitative evidence that nearly neutral mutations exist, there are few robust estimates how rapidly the proportion of effectively neutral mutations changes with effective population size.

We can potentially investigate how the proportion of effectively neutral mutations changes as a function of *N_e_* by considering how the number of non-synonymous (*p_N_*) relative to synonymous polymorphisms (*p_S_*) changes as a function of the number of synonymous polymorphisms (*p_S_*). The logic is as follows. Let us assume that synonymous mutations are neutral so that *p_S_* = *kN_e_U* where *k* is Watterson’s coefficient (=Sum 1/*i*), *N_e_* is the effective population size and *u* is the mutation rate. Let us further assume that non-synonymous mutations are either effectively neutral or deleterious so that *p_N_* = *kN_e_uf*, where *f* is the proportion of effectively neutral mutations and *k* is Watterson’s constant (Watterson 1975). We can therefore investigate how rapidly the proportion of effectively neutral mutations changes as *N_e_* increases by considering the correlation between *p_N_/p_S_* and *p_S_*. However, there are two issues. First, the two variables are not independent. We circumvent this problem by dividing *ps* into two halves using a hypergeometric distribution (Piganeau and Eyre-Walker 2009; James, et al. 2017); this means that *p_S1_* and *p_S2_* have uncorrelated sampling errors. We then consider the correlation between *p_N_/p_S1_* and *p_S2_*. Second, the theory outlined above assumes that the population is at equilibrium, however, non-equilibrium processes can potentially mimic the effects of variation in *N_e_*. Brandvain and Wright (Brandvain and Wright 2016) give an instructive example. Imagine a population, initially with no diversity. The population will accumulate genetic diversity through mutation, but the sites subject to deleterious mutation will approach their equilibrium frequency faster than neutral mutations because the equilibrium frequency is lower for these sites (Gordo and Dionisio 2005; Do, et al. 2015; Brandvain and Wright 2016). Hence, if we were to sample a locus through time in this population, we would observe a negative relationship between *p_N_/p_S_* and *p_S_*, despite the fact that *N_e_* has not changed since the population was founded.

The precise relationship between *p_N_/p_S_* and *p_S_* is potentially predictable. Ohta showed that the rate of evolution is proportional to 1/*N_e_* over a substantial range of *N_e_* values if the DFE is an exponential distribution (Ohta 1977). Kimura subsequently showed that under a gamma distribution with a shape parameter of 1/2 the rate was proportional to 1/*N*_*e*_^*0.5*^, which suggests, given that the exponential distribution is a gamma distribution with shape parameter of one, that the rate of evolution under a gamma distribution is proportional to 1/*N*_*e*_^β^, where *β* is the shape parameter. This general result was confirmed by Welch et al.(Welch, et al. 2008). Welch et al. (2008) also demonstrated that the number of polymorphisms and the nucleotide diversity are proportional to 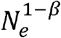; hence the nucleotide diversity at selected sites divided by the nucleotide diversity at neutral sites should be proportional to 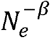 and hence there should be a linear relationship, with a slope of -*β* between the log of *p_N_/p_S_* and the log of *p_S_* if the DFE is a gamma distribution.

Several groups have shown that log(*p_N_/p_S_*) is negatively correlated to log(*p_S_*) across animal and plant species for nuclear DNA (Galtier 2016; Chen, et al. 2017) and across animal species for mtDNA (James, et al. 2017). It has also been demonstrated that this relationship holds across a single genome in *Capsella grandiflora, Arabidopsis lyrate* (Gossmann, et al. 2011) and the passenger pigeon (Murray, et al. 2017). This might be expected since the effective population size also varies across the genome in many species due to the effects of recombination, genetic hitch-hiking and background selection (Charlesworth 2009; Ellegren and Galtier 2016); regions of the genome with low rates of recombination, high mutations rates, gene densities or both, have lower effective population sizes than regions with high rates of recombination and a low density of selected mutations. However, no detailed study has been made within a species, and in particular, whether the relationship between log(*p_N_/p_S_*) and log(*p_S_*) is quantitively consistent with the nearly neutral theory of molecular evolution.

## Results

To investigate the relationship between the proportion of effectively neutral mutations and the effective population size, we compiled polymorphism data from 7918 autosomal genes from a North American population of *D. melanogaster*. We also use a smaller data set of 4676 autosomal genes with short introns (< 66 bp) as an alternative neutral reference (Halligan and Keightley 2006). Because we are interested in the relationship between log(*p_N_/p_S_*) and log(*p_S_*) and *p_N_* and *p_S_* can be zero for individual genes, we grouped genes into 10 groups of 791 genes (10 groups of 467 genes when using introns) and considered the correlation between log(sum(*p_N_*)/sum(*p_S1_*)) and log(sum(*p_S2_*)) (similar results were obtained with other groupings – see below).

As expected, given previous work, there is a positive correlation between *p_S_* and the rate of recombination (RR) (Spearman’s correlation r_s_ = 0.99, p<0.001) (Figure 1)(Begun and Aquadro 1992; Presgraves 2005; Langley, et al. 2012; Mackay, et al. 2012; Campos, et al. 2014). This positive correlation might be due to variation in the mutation rate or variation in *N_e_* across the genome. Since, there is no correlation between the synonymous site divergence (*d_S_*) between *D. melanogaster* and *D. yakuba* and RR (r_s_ = −0.10, p = 0.78), we conclude as others have done, that recombination is not mutagenic in Drosophila (Begun and Aquadro 1992; Betancourt and Presgraves 2002; Mackay, et al. 2012; Campos, et al. 2014), and that there is variation in *N_e_* due to either genetic hitch-hiking or background selection. The relationship between *p_S_* and RR is non-linear with a tendency for the number of synonymous polymorphisms to be substantially depressed in regions of low recombination (Figure 1).

**Figure 1.**
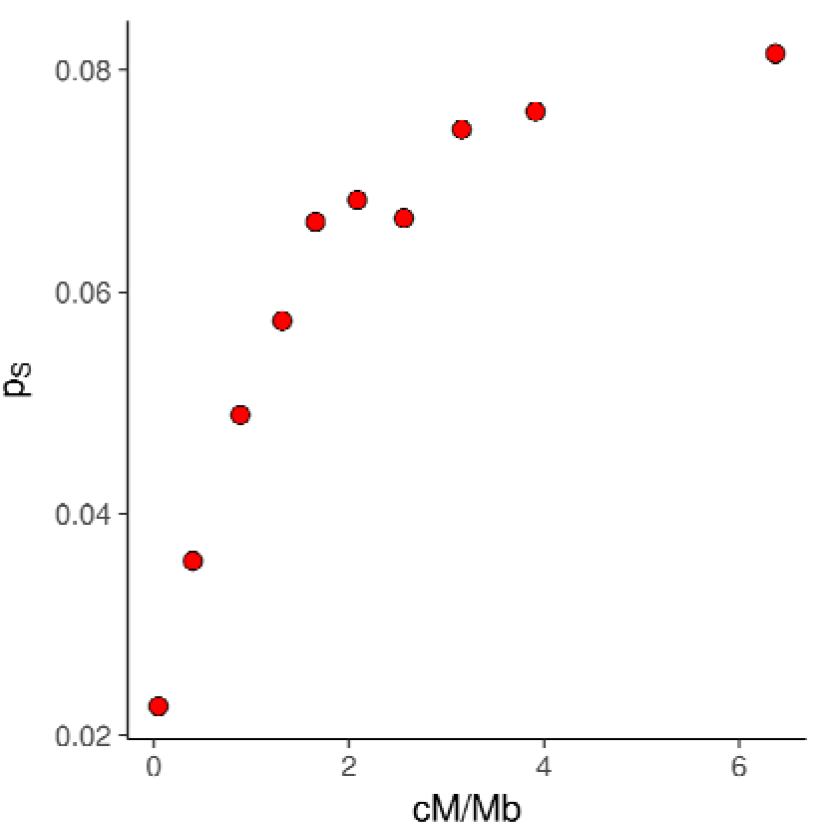
The correlation between the number of synonymous polymorphisms per site and the rate of recombination. Each point represents the average of 791 genes grouped according to their mutation rate.

We find that log(*p_N_p_S_*) is negatively correlated to log(*p_S_*) (slope = −0.53 (95% CIs = −0.48, −0.57), *p* < 0.001) (figure 2A). The relationship is almost perfectly linear, consistent with the underlying distribution of fitness effects being a gamma distribution. The slope is such that halving the effective population size increases the proportion of effectively neutral mutations by ~45%. The results are unaffected by using different grouping schemes (20 groups slope = −0.54 (−0.60, −0.48), 50 groups slope = −0.53 (−0.59, −0.48)).

**Figure 2.**
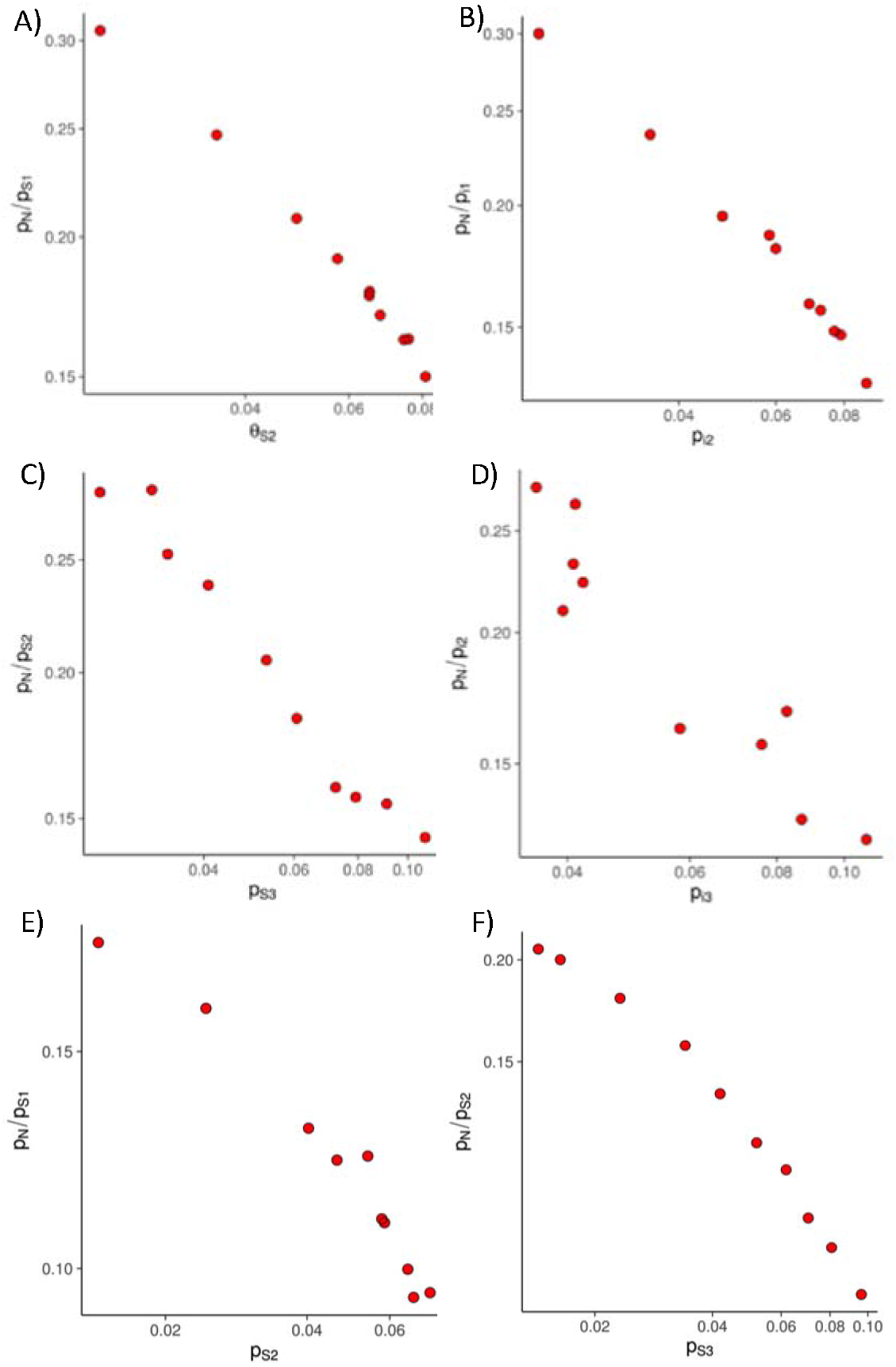
The relationship between *p_N_/p_S_* and *p_S_* or *p_i_*, grouping genes by either recombination rate, *p_S_* or *p_i_*. Panels A to D are using DGRP flies, panels E and F are using DPGP flies. A) using *p_S_* as the neutral standard, grouping genes by recombination rate, B) using *p_i_*, grouping genes by recombination rate, C) using *p_S_*, grouping genes by *p_s_*, D) using *p_i_* grouping genes by *p_i_*, E) using *p_S_*, grouping genes by recombination rate and F) using *p_S_*, grouping genes by *p_S_*.

We have assumed that synonymous mutations are neutral, whereas we know that there is selection on synonymous codon use in *Drosophila* species (Shields, et al. 1988; Akashi 1995). To investigate whether this affects our results we repeated the analysis using data from short introns as our neutral sites (Halligan and Keightley 2006). The relationship between log(*p_N_p_l1_*) and log(*p_12_*) is highly significant (*p* < 0.001) (figure 2B), almost perfectly linear and has a slope which is very similar to that found using synonymous sites (slope = −0.55 (−0.47, −0.65)). However, if there is selection on short introns then this would affect our conclusions.

In many species, we do not have a recombination rate map, so we also investigated the relationship between log(*p_N_/p_S_*) and log(*p_S_*) grouping genes by their *p_S_* or *p_l_* values rather than the RR. To do this we followed the method of James et al. (James, et al. 2017) in which *p_S_* (*p_l_*) is split three-ways using a hypergeometric distribution and *p_S1_* (*p_l1_*) is used to rank and group genes, *p_S2_* (*p_12_*) is used to measure *p_S_* (*p_l1_*) and *p_S3_* (*p_l3_*) is used to calculate *p_N_/p_S_* (*p_N_/p_l_*). The slopes using this method are −0.52 (−0.48, −0.56) and −0.64 (−0.53, −0.77) for synonymous and intron sites respectively (Figure 2C, 2D), confirming that this is a satisfactory method for investigating the relationship between log(*p_N_/p_S_*) and log(*p_S_*) when recombination rate data are not available and there is sufficient polymorphism data, although it should be noted that the results are subject to higher levels of sampling error.

We performed most of our analyses using the DGRP lines which were sampled from a derived population in the United States (Mackay, et al. 2012). To check that our results are consistent across populations we also performed the analysis on the DPGP2 lines sampled from Rwanda (Pool et al. 2012). Whether we group genes by RR or *p_S_* we observe a negative correlation between log(*p_N_/p_S_*) and log(*p_S_*) (Figure 2E, 2F) (grouping by *RR* slope = −0.42 (−0.34, −0.50); grouping by *p_S_* slope = −0.51 (−0.43, −0.58).

If the DFE is gamma distribution, then we expect the slope of the relationship between log(*p_N_/p_S_*) and log(*p_S_*) to be equal to the negative of the shape parameter of the DFE (Welch, et al. 2008). To investigate whether this is the case we estimated the shape parameter of the DFE using the site frequency spectrum (SFS) at non-synonymous and synonymous sites using the method of Keightley and Eyre-Walker (Keightley and Eyre-Walker 2007). We find that the shape parameter is significantly smaller (shape = 0.33 (0.30, 0.38)) than the slope of the relationship between log(*p_N_/p_S_*) and log(*p_S_*) (p < 0.001 bootstrapping the data by gene) for the American DGRP data and the African DPGP2 data (shape parameter = 0.35 (0.34, 0.36) which is significantly shallower than the slope, p<0.001). This discrepancy has been noted before across species for animal mtDNA (James, et al. 2017) and although not explicitly mentioned by the authors, is evident in the analysis between species in animal and plant nuclear DNA (Chen, et al. 2017).

There are a number of potential explanations for why the slope of the relationship between log(*p_N_/p_S_*) and log(*p_S_*) is greater than the shape parameter of the DFE estimated using the SFS. First, there could be problems with our method. However, previous simulations have suggested that the method recovers the expected relationship between log(*p_N_/p_S_*) and log(*p_S_*) under a gamma DFE in a model with background selection and linkage (James, et al. 2017).

Second, the pattern may be a product of the way in which the data have been processed. There three potential complications: inversions, admixture and identity-by-descent (IBD). Both datasets are known to have some admixture (Pool, et al. 2012; Pool 2015) and low recombination regions in the DGRP data show elevated admixture (Pool 2015), so our results could be affected, given the relationship between *p_S_* and RR. Similarly, inversion polymorphisms appear to increase levels of genetic diversity (Corbett-Detig and Hartl 2012; Pool, et al. 2012). We have removed inversions from the DGRP, and Campos et al. (Campos, et al. 2014) removed admixed regions from the DPGP data. In neither dataset were regions of IBD removed but these comprise a small fraction of the Rwandan dataset, in which this complication has been quantified – two individuals are estimated to be affected over 9 and 21 MB of their genome (Pool, et al. 2012). It therefore seems unlikely that these complications are responsible for our results given that the patterns we observe are consistent across two datasets which have different origins and have been processed differently.

Third, there might be a negative relationship between *N_e_* and the mutation rate. This seems unlikely within the *Drosophila* genome, given that there is no correlation between *d_S_*, a measure of the mutation rate, and the rate of recombination (see above). However, to investigate further, we split the synonymous (and intronic) divergence into two halves using the hypergeometric distribution then used *d_S1_* (*d_l1_*) to calculate the average *N_e_* for a group of genes (grouped by RR) as *p_S_/d_S1_* (*p_l_/d_l1_*) and used the other *d_S_* (*d_l_*) value as our estimate of the mutation rate. For synonymous sites, we find no correlation between our estimate of *N_e_* and *d_S_* (slope = −0.017, *p* = 0.54) but for introns we observe a positive and significant relationship (slope = 0.10, *p* < 0.001) (Supplementary Figures 1A and 1B); this latter correlation is consistent with the fact that *d_l_* is positively correlated to RR at the gene level (r_s_=0.09, p<0.001). These patterns are consistent with there being a positive correlation between RR and the mutation rate, a positive relationship which is offset at synonymous sites by greater efficiency of selection against deleterious mutations in highly recombining areas, leading to a lack of a correlation between synonymous divergence and RR. It therefore seems likely that there is a positive correlation between the mutation rate and *N_e_* in *Drosophila* and hence no evidence that they are negatively correlated. Similar results are obtained for the relationship between log(*p_N_/p_S_*) versus log(*p_S_/d_S_*) and log(*p_N_/p_i_*) versus log(*p_i_/d_i_*) grouping by RR (using synonymous SNPs, slope = - 0.50 (−0.48, −0.57); using intron SNPs, slope = −0.60 (−0.51, −0.70) or neutral polymorphism levels (−0.55 (−0.50, −0.59) and −0.54 (−0.47, −0.62)).

Fourth, the slope of the relationship between log(*p_N_/p_S_*) and log(*p_S_*) might be steeper than the estimate of the shape parameter of the DFE because the DFE is affected by *N_e_*; the shape parameter or the mean selection coefficient of new deleterious mutations might increase with increasing *N_e_*. To investigate this, we estimated the DFE using the SFS for each group of genes separately, grouping genes by the RR. The method estimates the shape parameter and the mean value of *N_e_s* for the DFE. Since we expect mean *N_e_s* to increase as *N_e_* increases, we divided our estimate of mean *N_e_s* by *p_S_* to yield an estimate of the mean strength of selection (equivalent results are obtained if *N_e_s* is divided by *p_S_/d_S_*). There is a marginally significant positive correlation between *p_S_* and the shape parameter (r = 0.72, p = 0.020) (Figure 3a), however the slope of this relationship is very shallow, and the shape parameter never approaches the (negative) of the slope of the relationship between log(*p_N_/p_S_*) and log(*p_S_*). The mean strength of selection is uncorrelated to recombination rate (r = 0.51, p = 0.13)(Figure 3b), but this is not a powerful analysis since the estimate of the mean value of *N_e_s* is subject to considerable uncertainty, even in a large sample such as the one we have used.

**Figure 3.**
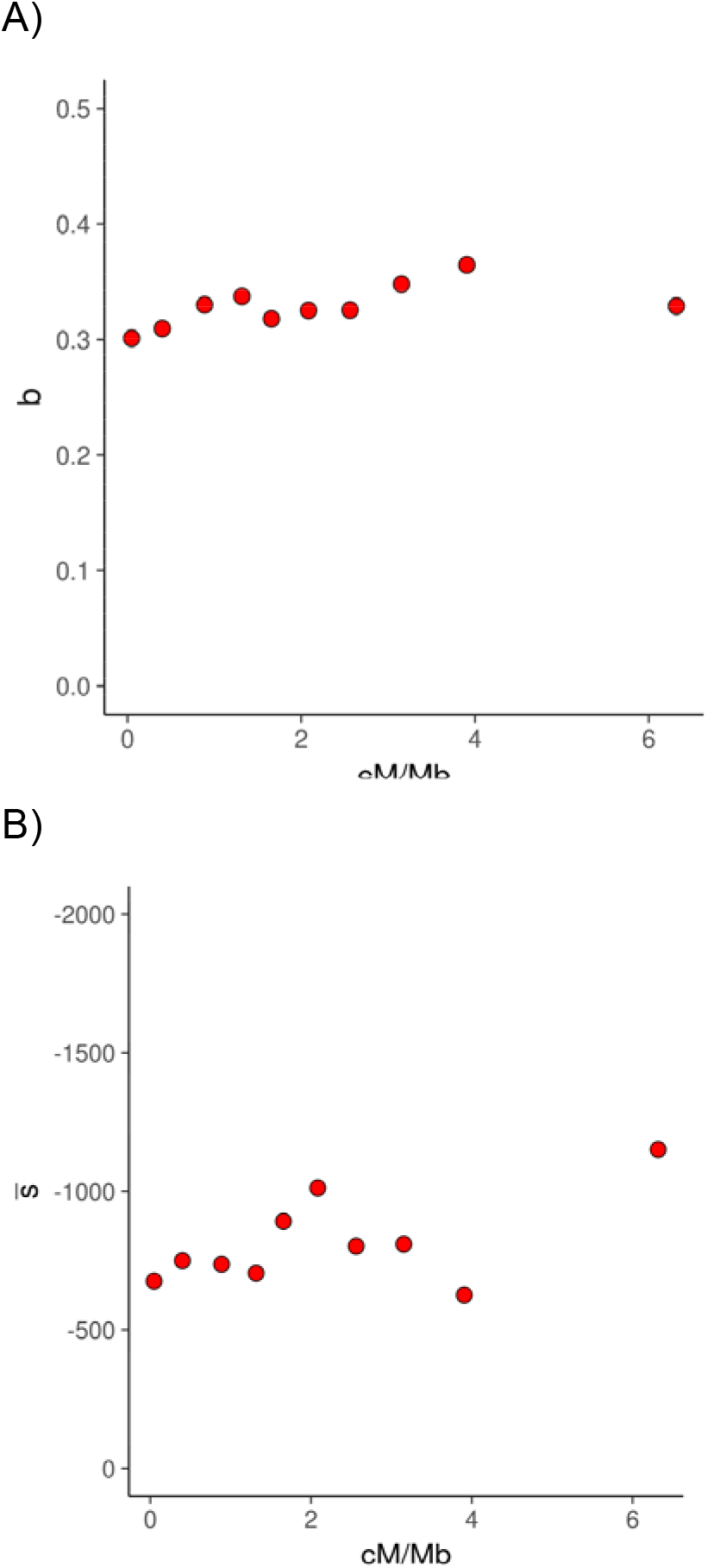
The relationship between the A) shape parameter and the B) estimated mean strength of selection and the rate of recombination.

Fifth, the difference between the slope of the relationship between log(*p_N_/p_S_*) and log(*p_S_*) and the shape parameter of the DFE estimated from the SFS might simply be due to the fact that the DFE is not well described by the gamma distribution and hence there is no expectation that the two quantities should be similar. The relationship between log(*p_N_/p_S_*) and log(*p_S_*) is very close to being linear for DGRP and DPGP data, suggesting that the gamma is a reasonable fit to the data. However, other distributions might give equally good fits.

Sixth, linked selection might affect the effective population size of neutral and deleterious mutations to different extents. We have previously shown, using simulation, that background selection does not affect the slope of the relationship between log(*p_N_/p_S_*) and log(*p_S_*) (James, et al. 2017), however hitch-hiking may affect the slope. It has been demonstrated, using simulated data, that deleterious genetic diversity recovers more rapidly after a bottleneck than neutral genetic diversity (Gordo and Dionisio 2005; Brandvain and Wright 2016) and this is also evident in HIV data after a hard-selective sweep (Pennings, et al. 2014).

To investigate whether hitch-hiking is responsible for the increase in the slope, we ran two sets of simulations. In the first, we simulated a nonrecombining locus composed of neutral sites and blocks of sites subject to varying levels of natural selection (i.e. the locus is composed of an equal number of sites subject to *Ns* = −1, −2, −4…etc). This locus was also subject to adaptive evolution, the rate of which was varied between simulations. We find, as expected, that nucleotide diversity for neutral and weakly selected sites decreases at the locus as we increase the rate of adaptive mutation (Figure 4a). However, the effect is attenuated for mildly deleterious mutations and there is no effect on diversity for strongly selected deleterious mutations, under the conditions of our simulation. As expected, the effect is attenuated if there is recombination between the advantageous locus and locus with neutral and deleterious mutations (Figure 4b, c, d). In these simulations we used a population size of 100, but similar results were obtained with 500 individuals (Figure S1).

**Figure 4.**
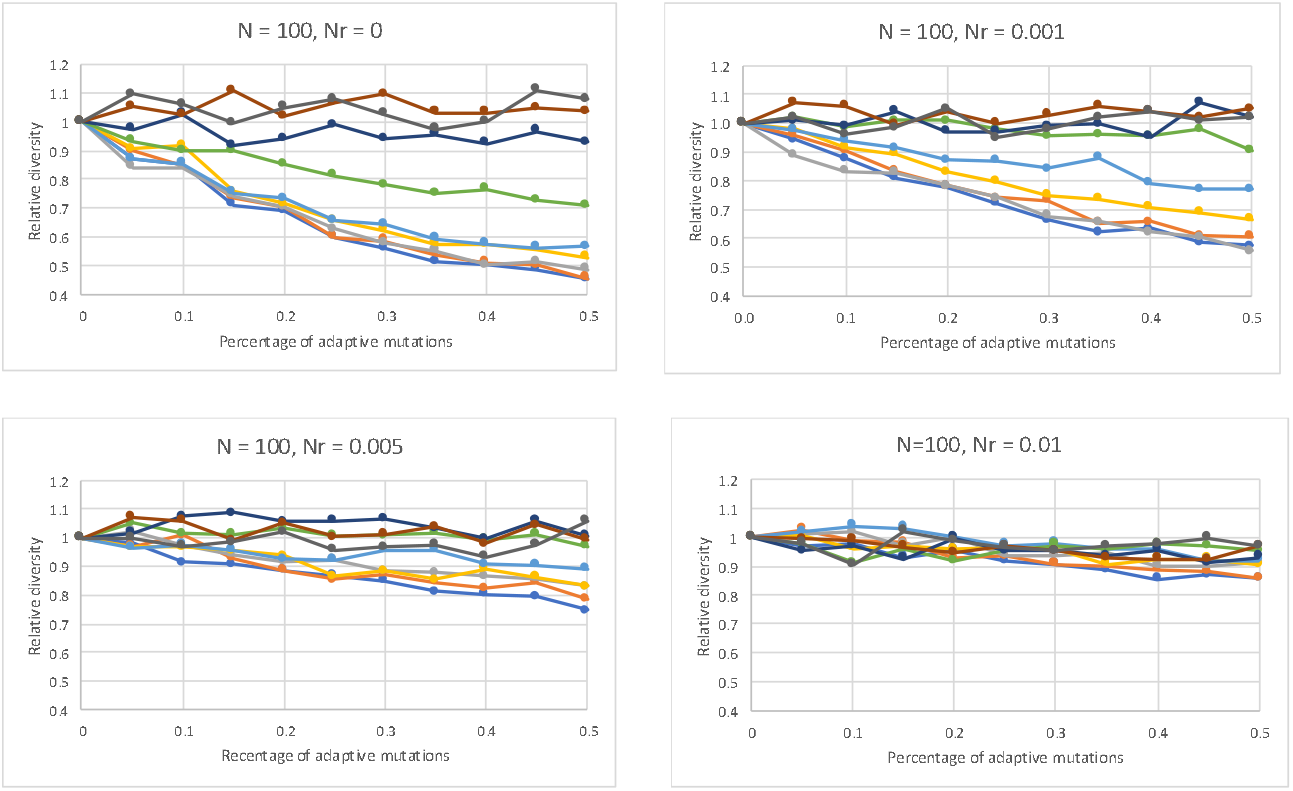
The effect of genetic hitch-hiking. The diversity at sites subject to different levels of negative selection, but all part of the same non-recombining locus, plotted against the frequency of advantageous mutation in the locus, for different population sizes and rates of recombination between the locus with advantageous mutations and that with deleterious and neutral mutations. *Ns* = 0 (dark blue), −1 (orange), −2 (grey), −4 (light orange), −8 (mid blue), −16 (green), −32 (dark blue), −64 (brown), −128 (black)

The fact that hitch-hiking depresses the diversity at neutral and weakly selected sites more than at strongly selected sites could potentially explain why the slope of log(*p_N_/p_S_*) and log(*p_S_*) is steeper than expected from the estimate of the shape parameter of the DFE; regions with high levels of hitchhiking will have low diversity, but the diversity of non-synonymous sites will be depressed less than that of synonymous sites; hence the slope between log(*p_N_/p_S_*) and log(*p_S_*) will be steeper. To investigate this, we ran a second set of simulations in which a locus, that experienced both neutral and deleterious mutations, whose fitness effects were drawn from a gamma distribution occurred, were linked to a locus undergoing adaptive evolution. For each rate of adaptive evolution, we averaged the number of neutral and selected polymorphisms per site across multiple sampling points and runs. We find the slope of log(*p_N_/p_S_*) and log(*p_S_*) is substantially and significantly steeper (slope = −0.67 (0.029)) than −0.4 (p<0.001), the negative of the shape parameter of the gamma distribution used to generate the deleterious mutations (Figure 5). Similar results were obtained using a population size of 500 (Figure S1); in this case the slope = −0.74 which is significantly greater than the expected value of −0.4 (p<0.001).

**Figure 5.**
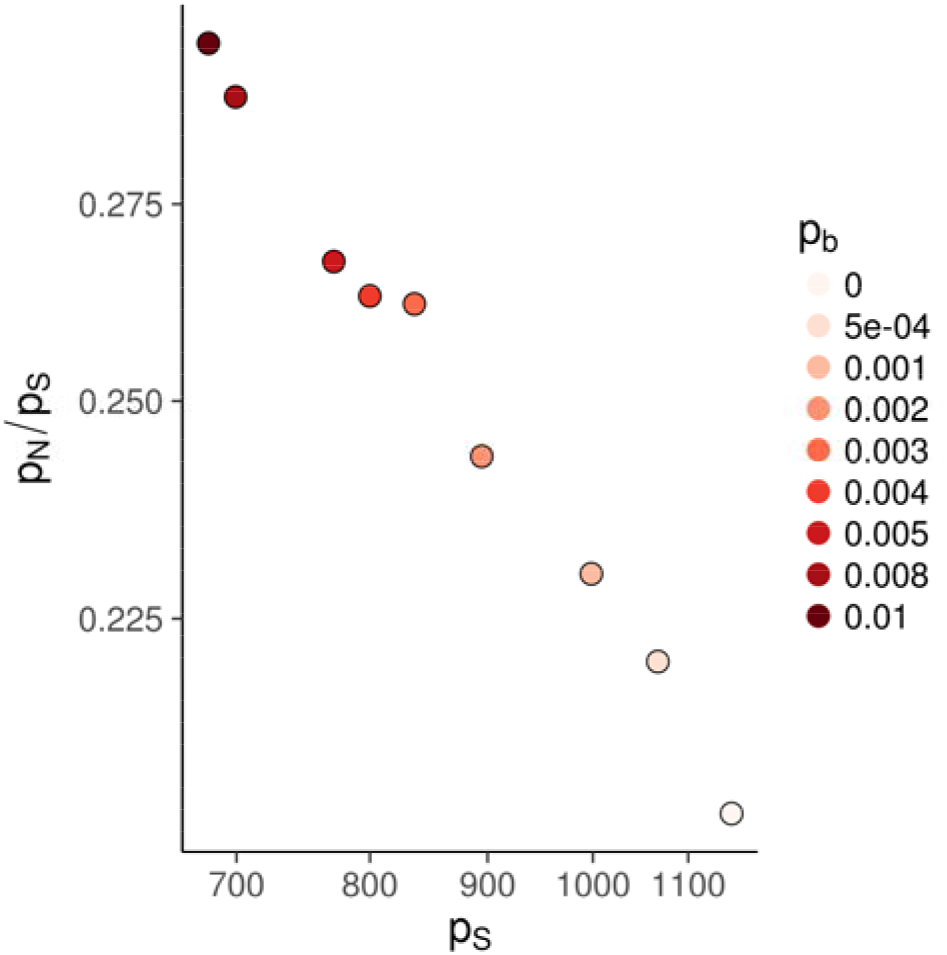
The effects of hitch-hiking. *p_N_/p_S_* plotted against *p_S_* both on log-scales, for simulations in which deleterious mutations are drawn from a gamma distribution for different rates of adaptive evolution. Each point represents the mean from a simulation run at a certain rate of beneficial mutation. The slope of the line, −0.67, is significantly steeper than the shape parameter of the gamma distribution used to simulate the data, which was 0.4.

## Discussion

We have shown that the proportion of mutations that are effectively neutral, as measured by the ratio of the number of non-synonymous and synonymous polymorphisms, decreases as the effective population size, as measured by the number of synonymous polymorphisms, increases across the *D. melanogaster* genome. On a log-log scale the relationship is approximately linear and the slope of the relationship is such that halving the effective population size increases the proportion of effectively neutral mutations by ~45%. The slope of the relationship between log(*p_N_/p_S_*) and log(*p_S_*) is similar to that found across plant and animal species for nuclear DNA (Galtier 2016; Chen, et al. 2017) and for animal mtDNA (James, et al. 2017).

Using data from across the *D. melanogaster* genome, we find, as others have between species (Chen, et al. 2017; James, et al. 2017), that the slope of the relationship between log(*p_N_/p_S_*) and log(*p_S_*) is steeper than we expect given an estimate of the DFE from the site frequency spectrum. We have previously shown by simulation that our method recovers the expected relationship when non-synonymous mutations are gamma distributed, even if there is substantial linkage between sites, so it seems unlikely that the steeper slope is due to a problem with our method (James, et al. 2017). We also find no evidence that there is a negative correlation between *N_e_* and the mutation rate across the genome or a change in the shape parameter or mean of the DFE. However, it is possible that the discrepancy is because the DFE is not gamma distributed. Although, the relationship between log(*p_N_/p_S_*) and log(*p_S_*) is approximately linear, as expected under the gamma distribution (Ohta 1977; Welch, et al. 2008), it is possible that other distributions, such as the log-normal would fit the data equally well. It is also possible that our estimate of the DFE is being influenced by mutations that are positively selected, particularly those subject to balancing selection. The influence of positively selected mutations on the estimation of the DFE and the relationship between log(*p_N_/p_S_*) and log(*p_S_*) is unknown.

We have shown that the steeper slope is consistent with a model of genetic hitch-hiking. We find, through simulation, that genetic hitch-hiking affects the relationship between log(*p_N_/p_S_*) and log(*p_S_*), making it steeper than expected. We demonstrate that hitch-hiking depresses the level of genetic diversity at neutral and slightly deleterious sites, but has little or no effect on the diversity of strongly deleterious mutations. As a consequence, regions of the genome with high rates of hitch-hiking have low neutral diversity, but deleterious genetic diversity is not as affected. This effect is likely due to the fact that genetic diversity recovers more rapidly at negatively selected sites than neutral sites after a bottleneck or hitch-hiking event (Gordo and Dionisio 2005; Pennings, et al. 2014; Brandvain and Wright 2016). The finding that hitchhiking can induce a relationship between log(*p_N_/p_S_*) and log(*p_S_*) raises the question of whether the relationship is due to variation in the efficiency of selection, non-equilibrium dynamics, or a combination of both. Resolving this question will require knowledge about the rates and strength of selection acting upon advantageous mutations and extensive simulation. The effect of hitch-hiking may also depend on demography – our simulation does not attempt to mimic the demographic changes that have affected *D. melanogaster*.

It is currently unclear how general these results are likely to be; are we likely to find a negative correlation between *p_N_/p_S_* and *p_S_* across other genomes? This will depend upon variation in the rate of recombination and density of selected sites establishing variation in *N_e_*, the distribution of fitness effects and/or variation in the rate of adaptive evolution across the genome. Gossman et al. (Gossmann, et al. 2011) and Murray et al. (Murray, et al. 2017) have recently shown that *p_N_/p_S_* is negatively correlated to *p_S_* across the two plant species and the passenger pigeon genome. Also many species appear to show similar levels of variation in *N_e_* across their genomes as *D. melanogaster* (Gossmann, et al. 2011) and hence might be expected to show a correlation between *p_N_/p_S_* and *p_S_*; initial work suggests that this is the case in both primates and rodents, but with some qualitative differences (Castellano and Eyre-Walker, unpublished results).

An open question is whether variation in *p_N_/p_S_* across a genome has any implications for phenotypic variation. Whether it does, depends on the genetic architecture of quantitative traits; is most of the variation due to common or rare mutations of large or small effect, and what role does natural selection play in shaping this architecture? It seems likely that most of the variation in fitness will be contributed by rare mutations of relatively large effect if genetic variation is maintained in mutation-selection balance (Eyre-Walker 2010). As a consequence, variation in *p_N_/p_S_* is unlikely to be important because neither variation in *N_e_* or non-equilibrium dynamics will affect the equilibrium frequencies of strongly deleterious mutations. Variation in *N_e_*, generated by for example background selection, has no effect, because the equilibrium frequencies of a deleterious mutation are independent of *N_e_*, and we have shown that hitch-hiking also has little effect on the diversity of strongly selected mutations. However, weakly selected mutations may contribute substantially to traits that are not subject to strong selection and hence variation in *p_N_/p_S_* across the genome may affect the architecture of these traits.

Our results suggest that the proportion of effectively neutral mutations varies across the *Drosophila* genome and declines as a function of synonymous diversity; this is likely to be due to two factors; variation in the effective population size and non-equilibrium dynamics. Interestingly, we show that genetic hitch-hiking has little or no effect on strongly deleterious genetic variation.

## Materials and methods

### Datasets

This study was carried out on the four large autosomes (2L, 2R, 3L and 3R) of *D. melanogaster*. The population genomic data came from two sources, Raleigh, North Carolina (DGRP) (Mackay, et al. 2012) and Gikongoro, Rwanda (DPGP)(Pool, et al. 2012). We used Freeze 1.0 *Drosophila melanogaster* Genetic Reference Panel (DGRP) project. Sites with residual heterozygosity and low-quality values were excluded from the analyses. We also excluded all regions which contain common inversion polymorphisms. The method to estimate the distribution of fitness effects of new mutations (DFE) requires all sites to have been sampled for the same number of chromosomes and since some sites were not successfully sampled in all samples we reduced the original data set to 128 chromosomes by randomly sampling the polymorphisms at each site without replacement. To estimate divergence from *D. yakuba* we randomly sampled one *D. melanogaster* chromosome per site.

Coding exon and short intron (≤65 bp) annotations from *D. melanogaster* were retrieved from FlyBase (release 5.50, http://flybase.org/, last accessed March 2013). Genes 1:1 orthologs across *D. yakuba* – *D. melanogaster* were obtained from FlyBase (http://flybase.org/). We obtained a multiple genome alignment between the DGRP isogenic lines (Mackay, et al. 2012) and the *D. yakuba* genome (Drosophila 12 Genomes, et al. 2007) using the BDGP 5 coordinates. This alignment is publicly available at http://popdrowser.uab.cat/ (Ramia, et al. 2012). For each gene, we took all non-overlapping coding exons, independently of their inclusion levels. When two exons overlapped, the largest was chosen for subsequent analyses. Only exons without frameshifts, gaps or early stop codons were retained. In this way, we tried to avoid potential alignment errors which would inflate our mutation rate estimates. Our final data set fulfilling all of these criteria contains 7,918 coding genes.

Exonic sequences were trimmed in order to contain only full codons. We define our sites “physically”, so we estimated the rates of substitution at sites of different degeneracy separately. Only zero-fold and 4-fold degenerate sites in exon core codons (as described by Warnecke and Hurst (Warnecke and Hurst 2007)) were used. To estimate the rate of synonymous substitutions, we restricted our analysis to those triplets coding the same amino acid in the two species (*D. melanogaster – D. yakuba*). In restricting our analysis to codons not exhibiting nonsynonymous differences we assume that the codon has undergone no amino acid substitution — this avoids having to compute the different pathways between two codons, which differ by more than one change and it is a reasonable assumption given the level of amino acid divergence. For 4-fold degenerate sites we used the method of Tamura (Tamura 1992) to correct for multiple hits; this method allows for unequal GC content and ts/tv bias. We calculated the number of substitutions and the folded site frequency spectrum (SFS) for 4-fold degenerate sites and zero-fold degenerate sites, using a custom Perl Script.

We used positions 8–30 of introns ≤65 bp in length as an alternative neutral reference for some analyses (Halligan and Keightley 2006). For intron sequences, the invariant GT and AG dinucleotides at the 5’ and 3’ splice junctions, respectively, are excluded before calculating divergence. Only genes with at least one short intron and with less than 10% of gaps in the aligned sequences were kept. 4,676 orthologous genes pass the intron quality criteria in our final data set. We use an *ad hoc* Perl Script to estimate the number of short intron substitutions and to compute the folded SFS. Multiple hits were corrected using the Jukes and Cantor method (Jukes and Cantor 1969).

Details of the assembly and data filtering of the Rwandan DPGP dataset can be found in Campos et al. (Campos, et al. 2012; Campos, et al. 2014). The dataset comprises 22 genomes sequenced from haploid eggs. They excluded regions that showed admixture with European populations (Pool, et al. 2012) but did not exclude tracts of identity-by-descent (IBD). The IBD tracts potentially affected two strains RG10 and RG15 and a total 30MB ((Pool, et al. 2012) supplementary Table 4); it is unlikely that such a small amount of IBD will affect our results. The numbers of synonymous and non-synonymous sites and polymorphisms and the SFS were estimated by Campos et al. (Campos, et al. 2014).

Recombination rates were taken from Comeron et al. (Comeron, et al. 2012). They estimated the rate of crossovers in 100 kb non-overlapping windows in cM/Mb units. The rate of crossing-over used for a gene is that of the rate in the 100kb window that overlapped the mid-point of the gene.

### Hypergeometric sampling

To investigate the correlation between *p_N_/p_S_* and *p_S_* ranking genes by its recombination rate we split *p_S_* into 2 independent variables (similar to the splitting done in Smith and Eyre-Walker (Smith and Eyre-Walker 2003); Piganeau and Eyre-Walker (Piganeau and Eyre-Walker 2009)). This was done by generating a random multivariate hypergeometric variable as follows:

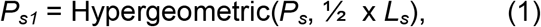

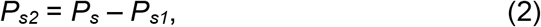

where *L_s_* is the total number of synonymous 4-fold degenerate sites and *P_s_* is the total number of 4-fold segregating sites. We divide Σ*P_s1_* and Σ*P_s2_* by ⅓ of Σ*L_s_* to get *p_S1_* and *p_S2_* at the bin level, respectively. We used *p_S1_* to estimate *p_N_/p_S1_* and *p_S2_* as a statistically independent estimate of the level of neutral diversity at the bin level.

To investigate the correlation between *p_N_/p_S_* and *p_S_* ranking genes by its level of neutral diversity we split *p_S_* into 3 independent variables following the scheme described by James et al. (James, et al. 2017). This was done by generating a random multivariate hypergeometric variable as follows:

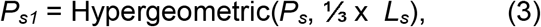

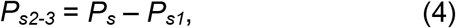

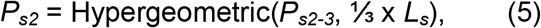

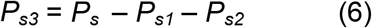

We divided Σ*P_s2_* and Σ*P_s3_* by ½ of Σ*L_S_* to get *p_S2_* and *p_S3_* at the bin level, respectively. We used *p_S1_* to rank genes and assign genes to bins, we then used *p_S2_* to estimate *p_N_/p_S1_* and *p_S3_* as a statistically independent estimate of the level of neutral diversity at the bin level.

### Gene binning strategy

To estimate the ratio between non-synonymous (*p_N_*) to synonymous polymorphisms (*p_S1_*) (or short intron, *p_i1_*) it is necessary to combine data from several genes because estimates from a single gene are noisy and often result in undefined values due to a lack of segregating synonymous (or short intron) sites. We therefore group genes into bins according to their rate of recombination or neutral polymorphism. Three different grouping schemes were used: 10 bins, 20 bins and 50 bins.

To investigate the relationship between the log of *p_N_/p_S1_* (or *p_N_/p_l1_*) and the log of our proxy for the *N_e_* ~ *p_S3_/d_S_* (or *p_l3_/d_l_*) we used the substitution rate at exon core 4-fold degenerate sites (or at short intron sites) as a proxy for the neutral mutation rate.

### Distribution of fitness effects estimated using the SFS

DFE-alpha (Eyre-Walker and Keightley 2009) models the DFE at functional sites by assuming a gamma distribution, specified by the mean strength of selection, *γ* = −*N_e_S*, and a shape parameter *β*, allowing the distribution to take on a variety of shapes ranging from leptokurtic to platykurtic. We ran DFE-alpha assuming a single instantaneous change in population size from an ancestral size *N_1_* to a present-day size *N_2_* having occurred *t_2_* generations ago using the folded SFS mode since the results are more robust to both demographic changes and linked selection (Messer and Petrov 2013). Provided the SFS at both neutral and functional sites, DFE-alpha infers *γ, β*, *N_2_/N_1_* and *t_2_*. We ran DFE-alpha (version 2.16) for each bootstrap replicate independently (see below) using the local version provided at: http://www.homepages.ed.ac.uk/pkeightl//software.

### Confidence intervals and p-value estimates

To calculate the 95% confidence intervals (CIs) for the slope (*b*) of the relationship between log(*p_N_/p_S_*) vs log(*p_S_*), we bootstrapped the data by gene 1,000 times. We split each bootstrap dataset into *N* bins (see Gene Binning Strategy above) and reestimated the slope for each replicate independently. To estimate the statistical significance of the difference between the slope (*b*) and the shape parameter (*β*) of the DFE using the SFS (Eyre-Walker and Keightley 2009) we estimated *β* and *b* for each bootstrap dataset. The p-value was the proportion of replicates in which −*b* > *β*.

### Statistical analyses

All statistical analyses are performed using the R statistical software (Team 2013). Linear regressions are carried out calling the R function “lm” (from the R package “base”). We calculate Spearman rank correlations (*ρ*) using the R function “cor.test” (from the R package “base”). The random hypergeometric variable is obtained through the R function “rhyper” (from the R package “stats”). All the code used to perform the analyses are available upon request from DC.

### Forward simulations

We used the forward simulation software SLiM, version 2.4.1 (Haller and Messer 2017) to simulate the behaviour of a single, 20 kb non-recombining locus with a mutation rate of 1×10^-6^, evolving over time in a small population of 100 individuals. Parameter values were chosen to generate substantial variation in the locus.

We ran two types of simulations. In the first, we investigated the impact of selective sweeps on different types of site, from strongly deleterious to neutral. Our locus was composed of 9 equally common categories of site at which mutations were subject to negative selection with 4*Ns* values of 0, −1, −2, −4, −8, −16, −32, −64, −128. We also allowed the locus to undergo adaptive evolution varying the total proportion of adaptive mutations from 0 to 0.5% We then investigated how the genetic diversity at each category of site changed as we increased the frequency of selective sweeps, such that adaptive mutations constituted between 0.05% and 0.5% of the total number of mutations that occurred in a simulation run. The adaptive mutations were subject to selection such that 4*Ns* = 400. For each simulation run, after an initial burn-in of 1000 generations we took samples of all mutations segregating in the population every 500 generations, corresponding to 5*N* generations so that samples should be largely independent. The simulations, were run for 25500 generations in total. For each set of parameters, we ran our simulations 30 times. We then averaged values of diversity over simulation runs. To compare the impact of hitch-hiking on sites under different levels of selection we divide the diversity for a particular set of sites (e.g. *Ns* = −4) by the mean diversity from the simulation with no adaptive evolution.

In the second set of simulations, the locus was divided into two, with half of the mutations being neutral and the other half deleterious mutations which were sampled from a gamma distribution with a shape parameter 0.4 and a mean s of 10, which is roughly in line with what we observe in estimates of the DFE from real data. As before, we altered the frequency at which a locus experienced selective sweeps by changing the frequency with which adaptive mutations occurred. We used 12 different frequencies of adaptive mutations, such that adaptive mutations constituted between 0 and 1% of the total mutations that occurred in a simulation run. We used the same burn-in and sampling procedure as in our previous set of simulations, and again for each set of parameters we ran our simulations 30 times. We then calculated average values of *p_S_* and *p_N_/p_S_* across our simulation runs. Code to run and analyse the forward simulations can be found at https://figshare.com/articles/Code_for_producing_the_simulations_in_Nearly_Neutral_Evolution_Across_the_Drosophila_melanogaster_Genome_/3084202

## Acknowledgements

We are grateful to Meg Woolfit for early work on this project many years ago and for Matt Webster for reinvigorating our interest in this problem. Funding for this study was provided by the University of Sussex and NERC, grant number NE/L502042/1.

---

**Table S1.**
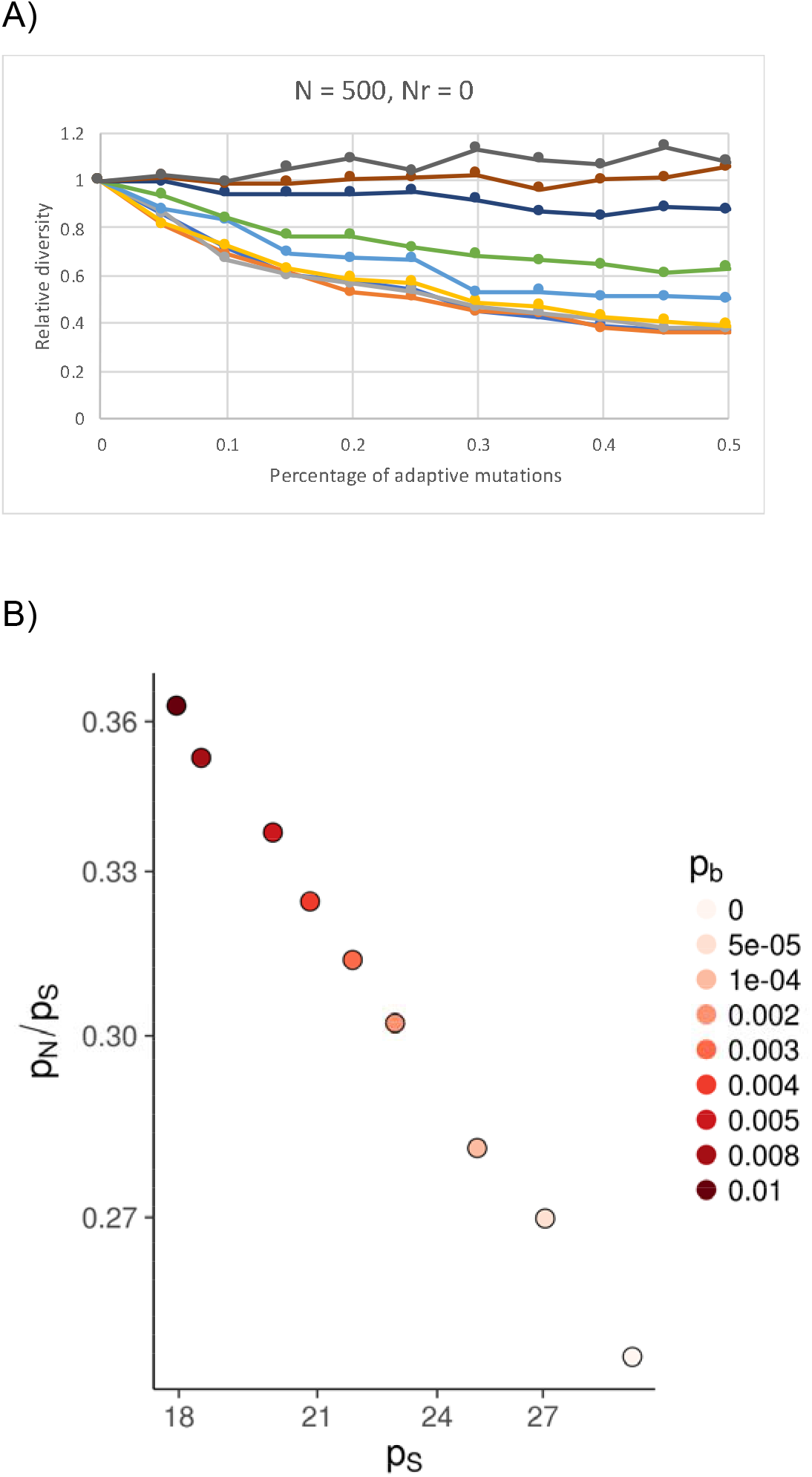
The influence of population size on simulation data. We repeated two sets of simulations with a population size of 500, rather than 100. A) Relative diversity versus the percentage of mutations that were advantageous in a non-recombining locus with equal number of sites of *Ns* = 0 (dark blue), −1 (orange), −2 (grey), −4 (light orange), −8 (mid blue), −16 (green), −32 (dark blue), −64 (brown), −128 (black). B) p_N_/p_S_ versus p_S_ for simulations in which mutations were drawn from a gamma distribution.

